# Rehabilitative Experience Interacts With FGF-2 to Facilitate Functional Improvement After Motor Cortex Injury

**DOI:** 10.1101/2025.10.02.680152

**Authors:** Alane Witt-Lajeunesse, Jan Cioe, Bryan Kolb

## Abstract

This experiment compared the effects of a structured rehabilitation regime (skilled reaching for 5 mo) and more varied training (complex environment for 5 mo) with and without postlesion infusion of FGF-2 for 7 days in rats having unilateral motor cortex lesions. Animals were tested on a motor battery throughout the five-month recovery period. The structured rehabilitation alone was ineffective in improving function whereas complex housing did improve performance on several measures. FGF-2 alone was ineffective but in combination with either the rehabilitation training or complex housing it did provide functional benefit. The combination of complex housing and FGF-2 was most effective as all motor measures showed significant improvement. Golgi analysis of layer III cortical pyramidal neurons showed that the complex housing essentially reversed the dendritic loss in the lesion animals. Curiously, there was no effect of FGF-2 on the cells measured, even though there was a beneficial effect of the combined FGF-2 and complex housing. It appears that varied rehabilitative programs, in combination with factors that promote neuronal plasticity are far more beneficial than similar training alone.

---

Various forms of rehabilitative training have been used to provide some benefit after focal cortical injury in both humans and laboratory animals (e.g., Biernaskie, Chernenko, & Corbett, 2004; Johansson, 2004; Johansson & Belichenko, 2002; Jones, Chu, Grande, & Gregory, 1999; Kleim, Jones, & Schallert, 2003; Kolb, 1995). Although the mechanisms of action of most treatments are poorly understood, it is presumed that the treatments somehow facilitate reparative processes in compromised tissue or stimulate the organization of compensatory networks that can facilitate functional improvement (for a review see Cramer et al., 2011). One likely mechanism, however, is the enhanced expression of neurotrophic factors such as Brain Derived Neurotrophic Factor (BDNF), Nerve Growth Factor (NGF), and basic Fibroblast Growth Factor (FGF-2) (e.g., Dahlqvist et al., 1999; Johansson, 2000). Although direct administration of these factors has been used as a treatment for brain injury (e.g., Kawamanta et al., 1997; Kolb, Cote, Ribeiro-da-Silva, & Cuello, 1997; Schabitz et al., 2004), the functional improvement is generally far from complete. The interplay between behavioral therapies and neurotrophic factors has not been well examined to date, however. Thus, the purpose of this study was to examine the benefits of the combination of either a structured therapy (repetitive training in a single motor task), or a more unstructured therapy (complex housing), and the administration of FGF-2. We chose to use FGF-2 because its expression is increased by complex housing and that administration of FGF-2 to animals with perinatal cortical lesions produced large improvements in functional outcome (e.g., Comeau et al., 2008; Gibb & Kolb, 2022; Monfils et al., 2006; 2008; Nemati & Kolb, 2011; Seo et al., 2013; Tung et al., 2017). Rehabilitation-induced recovery of function was assessed at specified intervals over 4 months using a battery of tests sensitive to motor cortex damage that we had previously found to be sensitive to focal motor cortex injury (Gonzalez & Kolb, 2003). The assessment protocol included a reaching task, spontaneous vertical exploration task, forepaw inhibition during swimming, tongue extension, single pellet reaching, and claw cutting. Assessment began three weeks after surgery, which is about the time that most spontaneous recovery has plateaued (Whishaw, Alaverdashvili, & Kolb, 2008). The complexity of dendritic fields in pyramidal neurons in cortex adjacent to the lesions was measured postmortem using Golgi-Cox procedures. We chose these cells because we had previously found them to be responsive to NGF and to complex housing and because they show significant increases in dendritic length following perinatal FGF-2 treatments (e.g., Gibb & Kolb, 2022; Kolb et al., 1997; Kolb et al., 2003).

## Methods and Procedures

### Animals

Subjects were 54 adult male rats, 400-675 grams, from the Charles River Long-Evans strain. Animals were randomly assigned to one of three groups: control, unilateral motor cortex lesion, and unilateral motor cortex lesion with post-lesion infusion of FGF-2. Treatment groups consisted of no treatment (*n* = 18), reaching training (*n* = 18), or complex housing (*n* = 18). No treatment and reaching training groups were housed in hanging cages in groups of four. Complex housing groups were placed in groups of five or six in complex cage housing [1.8 m high x 1 m deep x 1.5 m wide] containing runways, platforms, rope, trampoline, and small hanging cages attached to the wire mesh front.

All animals were maintained on a 12 hr light / 12 hr dark cycle at a temperature of 22° C. They were given ad-lib food and water, except during training and testing, when they were food-deprived to 85% of their body weight. Animals receiving complex cage housing were placed there several days prior to receiving surgery to get accustomed to their environment. Animals were maintained according to the regulations of the Canadian Council on Animal Care.

### Pre-training

Prior to being placed into treatment groups, all rats were pre-trained on a tray reaching task (Whishaw, O’Connor, & Dunnett, 1986; Whishaw, Pellis, Gorny, & Pellis, 1991). The preferred paw was considered the dominant paw for testing protocols. Rats were trained and tested in nine Plexiglas cages measuring 28 cm deep x 19 cm wide x 26 cm high with clear tops. The fronts were constructed of 2-mm bars separated from each other by a 9-mm gap. A 4-cm wide and 0.5-cm deep tray containing chick feed was mounted on the front of each cage. The rat had to reach through the bar, grasp the food, and then retract it into the box. If the rat dropped the food, it fell through the mesh floor and was lost. Rats were trained 20 min per day until they all learned the task (approximately 2 weeks). After learning the task, they were then videotaped for 5 minutes and their reaching was scored using a frame-by-frame analysis. A *reach* was recorded when the rat touched the food but had not been able to put into its mouth; a *hit* was counted when the rat reached for the food and ate it. Percentages were determined for each paw, and paw dominance was established based on these data.

### Surgical Procedures

Lesion and lesion+FGF-2 rats were anesthetized with injections (ip) of sodium pentobarbital (65 mg/kg) and atropine methyl nitrate (5 mg/kg). Surgeries were performed according to procedures described by Rowntree and Kolb (1997) and Kolb, Cioe, and Whishaw (2000b). Craniotomy was performed by removing the bone over the motor cortex from 1 mm lateral to the midline to the sagittal ridge (4 mm lateral) and from -4 mm bregma anterior to +2 mm bregma. After craniotomy and incision of the dura, focal unilateral suction lesions (6 mm long x 3 mm wide) were made. These aspiration lesions included Zilles’ (1985) areas Fr1, the lateral part of Fr2, the posterior part of Fr3, and area FL. Lesions were made in the motor cortex contralateral to the dominant paw.

Animals were given FGF-2 according to similar procedures described by Kolb et al., (1997). The animals received intracerebral ventricular infusions of 1 µg human recombinant FGF-2 (or vehicle) for 7 days via a minipump placed into the intact hemisphere immediately after lesions were performed. Stainless steel (23 gauge) cannulae were implanted into the intact lateral ventricle at the following coordinates relative to bregma: anterior/posterior, -0.8 mm; lateral, 1.5 mm; and ventral, 3.5 mm from the skull. The minipumps and tubing were removed from anaesthetized animals 1 week after implantation.

### Behavioral Treatment

Animals were placed into one of three treatment groups: no treatment (NT), reaching training (RT), and complex housing (CH). Animals who were in the NT groups stayed in their home cages and were handled when cages were cleaned or during testing. Reach training consisted of daily skilled reaching training for 1 hr on the tray reaching task, 5 days per week over a period of 4 months. Training began 3 days after injury while the animals were still receiving the FGF. This type of training was considered a structured treatment because there were specific muscle groups being utilized for reaching. Complex housing consisted of rats living 24 hr per day in the complex housing environment. This environment encouraged the use of a variety of muscle groups through the ability to move in an enlarged area with additional stimuli to encourage interaction. During training and testing some animals in the RT and CH groups were given bracelets to prevent reaching with the ipsilateral or non-dominant paw if necessary. A strip of elastic plaster [2 cm wide x 6 cm long] was wrapped around the rat’s forearm prevent the animal from using its non-dominant paw in reaching through the bars to retrieve food. Most of the rats learned to use their dominant (impaired) limb even when the bracelets were not present (Whishaw, 2000).

### Behavioral Testing

Testing took place every 14-18 days over 4 months for all groups. Rats were videotaped for all tests. The battery of tests included the following: (a) a skilled reaching test for 5 min, (b) a spontaneous vertical exploration test for 5 min, (c) forepaw use during swimming, (d) single pellet reaching, and (e) tongue extension. Brain weights, cortical thickness, and claw cutting abilities were examined after the rats were sacrificed.

### Skilled reaching

This test was the same as the skilled reaching task described previously. Scoring consisted of taking the total number of successful *hits* divided by the total number of *hits* and *reaches* x 100 for each paw to establish a percentage. The paw with the highest percentage of hits was considered the dominant or preferred paw.

### Spontaneous vertical exploration

This test is sensitive to chronic limb use asymmetries (Jones & Schallert, 1994; Liu et al., 1999; Schallert & Lindner, 1990). Animals were individually placed in a clear Plexiglas cylinder (30 cm in diameter & 45 cm high) for 5 min. Rats freely explored the space by using their forelimbs on the cylinder wall. Vanilla extract or chocolate chip mash was dabbed around the top of the cylinder to motivate the animals to explore the wall. The cylinder was placed on a glass table with a mirror angled at 45° below the glass so that the forelimbs could be viewed. Two behaviors were scored: (a) independent use of the left or right forelimb contacting the cylinder wall during rearing, initiating of a weight-shifting movement, or moving laterally along the wall in a vertical position and (b) simultaneous use of both the left and right forelimbs (within 0.5 s of each other) during a rear or a move laterally along the wall. Scoring consisted of establishing a percentage of non-dominant paw contacts divided by the total number of dominant, non-dominant, and bilateral wall contacts x 100 to establish a percentage. Intact animals tend to use one paw or both paws approximately 50% of the time, whereas unilateral lesion animals tend to use the non-dominant paw (i.e., the paw ipsilateral to the lesion).

### Forepaw use during swimming

When swimming forward in water, normal rats tend to hold both forepaws under their chin, keeping the paws immobile and using their hindlimbs to propel themselves (Schapiro, Salas, & Vukovich, 1970). In unilateral lesion rats that have learned to swim to a visible platform, only the unaffected forelimb tends to remain immobile, while the affected forelimb produces strokes along with the hindlimbs (Stoltz, Humm, & Schallert, 1999). Rats were tested in a rectangular aquarium (90 cm x 43 cm x 50 cm) with methods similar to Stoltz et al. A visible 9-cm^2^ square wire mesh platform was at one end of the aquarium; water temperature was maintained at approximately 22° C. During the training phase, animals were released from the opposite end of the tank and consecutive trials were given until they swam directly to the platform without touching the sides of the tank. After the animals completed five successful trials, they were given pieces of chocolate chip cookies or food pellets and placed under a heat lamp as reinforcement. A swim score was quantified by summing the number of strokes made with the impaired forelimb minus the number of strokes made with the unimpaired forelimb as a mean of all five trials. This was considered the *forepaw inhibition index*.

### Tongue extension

Tongue protrusion in rats can be examined to determine whether feeding abnormalities are present in lesion animals (Whishaw & Kolb, 1983). To examine tongue extension, a spatula covered with a mash of chocolate chip cookies and warm water was placed flush to the front of the home cage at right angles to the bars where it would be easiest for the animals to reach; each animal was tested individually. The area that the animal licked was then measured.

### Single pellet reaching

Rats were tested in single pellet reaching boxes to examine qualitative features of the actual reaching movements (as described in Metz & Whishaw, 2000; Whishaw, 2000; Whishaw, Pellis, Gorny, Kolb, & Tetzlaff, 1993). Boxes were made of clear Plexiglas 11 x 38 x 40 cm high. In the exact center of the front wall was a 1.5 cm wide slit that extended from the floor to a height of 31 cm. On the outside of the wall, in front of the slit, mounted 4 cm above the floor, was a 3-cm wide by 10.5-cm long shelf. Two small indentations, each .5 cm in diameter, were located on the floor of the shelf. These held food pellets. The indentations were 2 cm away from the inside wall of the box and were positioned on the outer edges of the slot. Food was placed in the indentation contralateral to the dominant paw for each rat. Training involved having the animal successfully reach for and receive a food pellet. After a successful reach, a food pellet was placed in the back of the box to encourage the rat to move from the front opening and reposition itself prior to the delivery of the next food pellet.

Three successful reaches for each rat were rated for qualitative features of the movement. These movements included the following: (a) *orient*, the head is oriented to the target food and sniffing occurs; (b) *limb lift*, the body weight is shifted to the back, the limb is raised from the floor with the upper arm, and digits are moving to the midline; (c) *digits close*, the digits are semi-flexed and the paw is supinated with the palm facing toward midline; (d) *aim*, the upper arm raises the elbow, adducting it so that the forearm is midline with the body and the paw is under the mouth; (f) *advance*, the head is lifted, the elbow is adducted, and the limb is directed to the target as the body weight shifts forward and laterally; (g) *digits open*, the digits are extended as the limb moves forward; (h) *pronation*, the upper arm moves, abducting the elbow and the paw moves directly over the food with the palm down; (i) *grasp*, the arm is still as the digits close onto the food, then the paw lifts; (j) *supination I*, the limb withdraws, the elbow is adducted, and the paw is supinated 90°, facing medially before leaving the slot; (k) *supination II*, the head is pointed downward, the paw is supinated another 90°, as food is presented to the mouth; and (l) *release*, the digits are opened and food placed into the mouth. Each of the movements was rated on a 3-point scale. If the movement appeared normal, it was given a score of 0; if the movement appeared slightly abnormal but was recognizable, it was given a score of 1; and if the movement was absent or compensated entirely by movement of a different body part, a score of 2 was given. Animals were videotaped until three successful reaches were made and scores were established for each of the 11 reaching components. A successful reach was one in which the animal reached for and ate the pellet on the first attempt. An independent scorer rated the animals.

### Claw cutting

Examination of claw length can provide information about an animal’s behavioral competencies. Loss of the ability to claw cut appears to be due to inefficient biting and chewing as opposed to grooming (Whishaw, Kolb, Sutherland, & Becker, 1983). Claws were examined according to procedures established by Whishaw et al. (1983). When the rats were sacrificed for brain histology, the length of all the rear paw claws was measured to the nearest .5 mm. The measure was made from the cuticle (tissue surrounding the proximal edge of the claw) to the tip. Mean claw length for each paw was recorded.

### Preparation of Brains

After all testing was completed, rats were deeply anesthetized with .5 cc of euthanol and intracardially perfused with a solution of 0.9% physiological saline. The brains were extracted whole and weighed. To weigh the brains, the olfactory bulbs were blocked off 2 mm from the tip of the cerebral hemispheres, and the cerebellar flocculi and pineal gland removed. Brains were then placed into 20 ml of Golgi-Cox solution (Glaser & van der Loos, 1981). Brains remained in this solution for 14 days and then were placed into 30% sucrose for at least 3 days prior to being sectioned. Brains were photographed, blocked perpendicular to the midline at approximately the level of the anterior commissure, and blocked again through the caudal portion of the occipital cortices. Coronal sections were cut in 200 µm on a vibratome into 6% sucrose solution, mounted on 2% gelatin-coated slides, and developed according to methods by Gibb and Kolb (1998). Cortical thickness was measured according to methods described by Stewart and Kolb (1988). Golgi-Cox stained sections were projected on a Zeiss DL 2 POL petrographic projector set at a magnification of 10x. Measurements made with a plastic millimeter ruler were taken at three points in each of five planes. Plane 1: First plane with caudate-putamen visible (Zilles’ areas Gu, Par 1, Fr 2). Plane 2: Anterior commissure (Zilles’ areas Par 2, Par 1, Fr 1). Plane 3: First hippocampal section (Zilles’ areas Gu, Par 1, Fr 1). Plane 4: Posterior commissure (Zilles’ areas Te 1, Oc 2L, RSA). Plane 5: Most posterior hippocampal section (Zilles’ areas Te 1, Oc 1B, Oc 2ML). All measurements were made without knowledge of the group identity of the animals by an independent examiner.

### Golgi analysis

Layer III pyramidal cells in Zilles’ area Par 1 were traced using a camera lucida at 250X. In order to be included in the data analysis, the dendritic trees of pyramidal cells had to fulfill the following criteria: (a) the cell had to be well impregnated and not obscured with blood vessels, astrocytes, or heavy clusters of dendrites from other cells; (b) the apical and basilar arborizations had to appear to be largely intact and visible in the plane of section. The cells were drawn and analyzed using a Sholl analysis for estimation of dendritic length (Sholl, 1956) was performed. For this analysis a transparent overlay of concentric circles spaced 20 µm apart was placed over the neuron drawing by centering the innermost ring in the middle of the cell body. The number of dendrite-ring intersections was counted for each ring and the total number used to estimate total dendritic length in µm (number of intersections X 20). Five cells were drawn in each hemisphere of each rat. The statistical analyses were done by taking the mean of the measurements on the five cells for each hemisphere of each subject. Analyses were done separately on the two hemispheres, one ipsilateral and one contralateral to the preferred forepaw.

The cells in Par 1 were chosen because these cells were next to the lesion area, and we had shown previously (Rowntree & Kolb, 1997) that these cells showed atrophy after similar lesions and that treatment with neutralizing antibodies enhanced the atrophy and this increased atrophy was correlated with poorer functional outcome. Furthermore, cells in Par 1 are especially sensitive to the effects of complex housing and we had shown that complex housing-induced changes in these cells were associated with recovery after cortical injury (Kolb & Gibb, 1991).

### Statistical Methods

We were interested in two primary measures: (a) improvement over time as assessed by repeated measures analyses of variance (ANOVAs) and (b) improvement at the end of training as assessed by doing a two-way ANOVA with lesion group and training as factors on test day 5. Fisher’s PLSDs were used for post hoc evaluations.

### Anatomical Results

The lesions removed the intended tissue and were consistent across treatment groups as illustrated in Figures 1 and 2. There was white matter damage in all lesion animals as the external capsule degenerated under the lesion. There was no obvious FGF-2 cell proliferation either in the subventricular zone or in the perilesional region.

**Figure 1.**
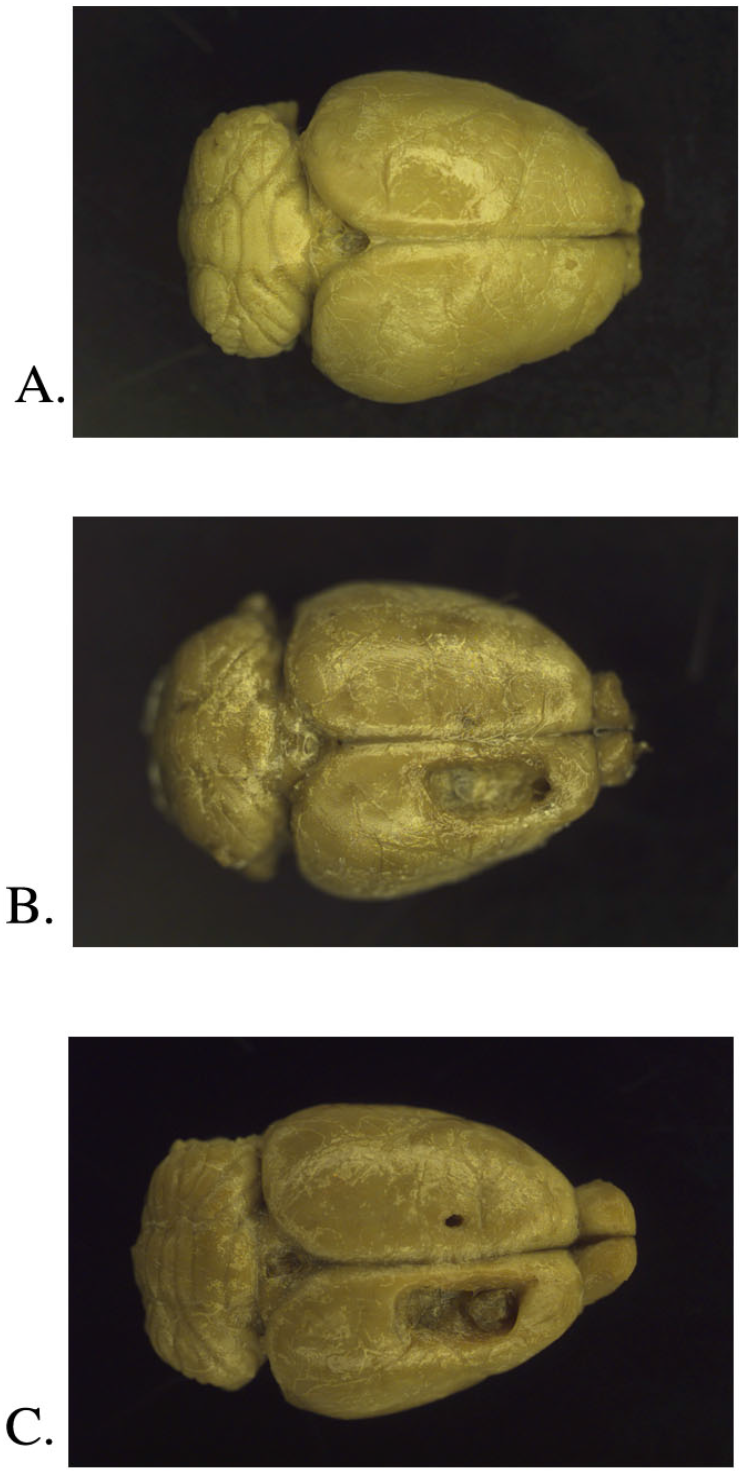
Representative examples of control (A), lesion (B), and FGF-2 (C) brains.

**Figure 2.**
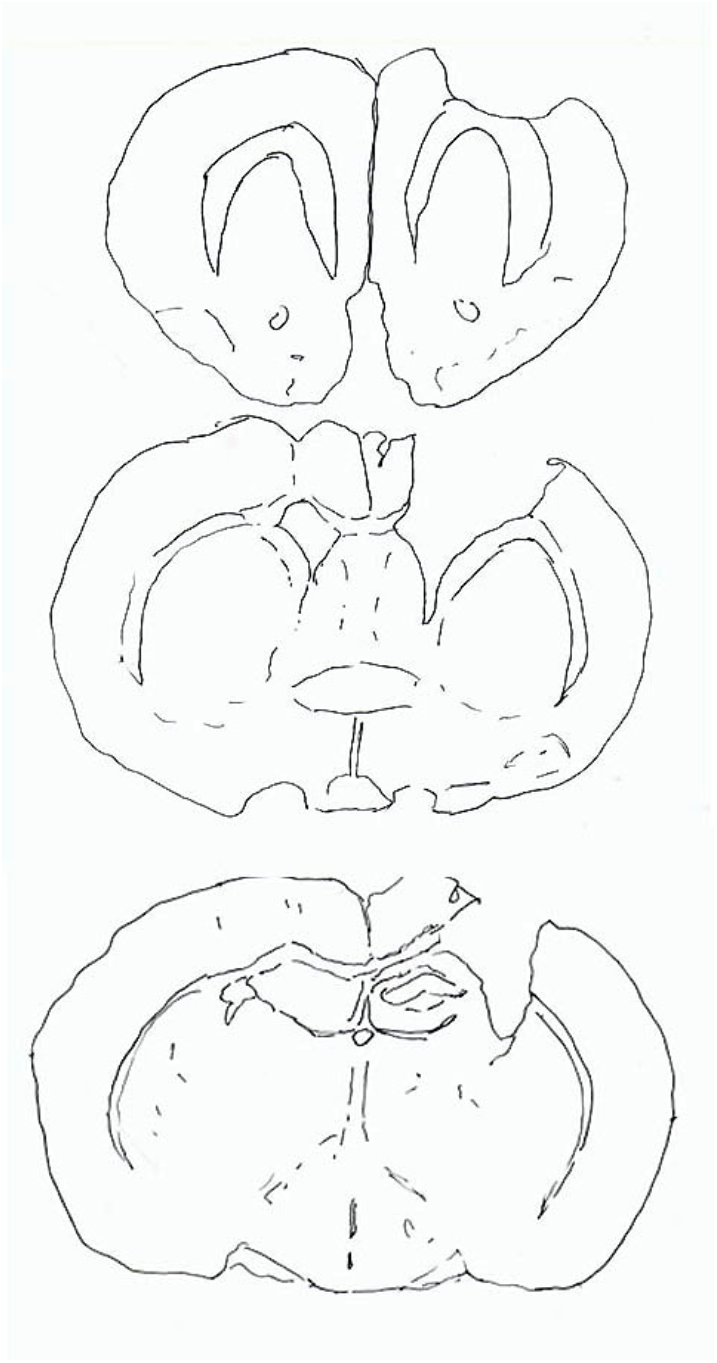
Serial drawing of Golgi-Cox stained-coronal sections through the brain of a representative unilateral motor cortex lesion rat.

### Cortical Thickness

Cortical thickness was analyzed according to lesion (dominant) or intact (non-dominant) hemisphere. Cortical thickness in control animals was determined by using the hemisphere contralateral to the dominant paw. Overall, the lesion and lesion+FGF-2 groups had thinner cortices in the lesion hemisphere, as can be seen in Figure 2 and 3. A two-way ANOVA was performed on the mean cortical thickness measurements averaged across all planes with lesion group and treatment as factors. In the lesion hemisphere there was a group effect, *F*(2, 37) = 12.44, *p* < .0001, but no treatment effect, *F*(2, 37) = .27, *p* = .764, nor interaction, *F*(4, 37) = 2.22, *p* = .0856 (Figure 3). Interestingly, the ANOVA for the intact hemisphere showed no group effect, *F*(2, 37) = .50, *p* = .610, no treatment effect *F*(2, 37) = 1.01, *p* = .375, but an interaction, *F*(4, 37) = 3.26, *p* = .017. The interaction had to do with a larger cortical thickness in plane 1 in the lesion+FGF-2 groups, mainly in the complex housing group.

**Figure 3.**
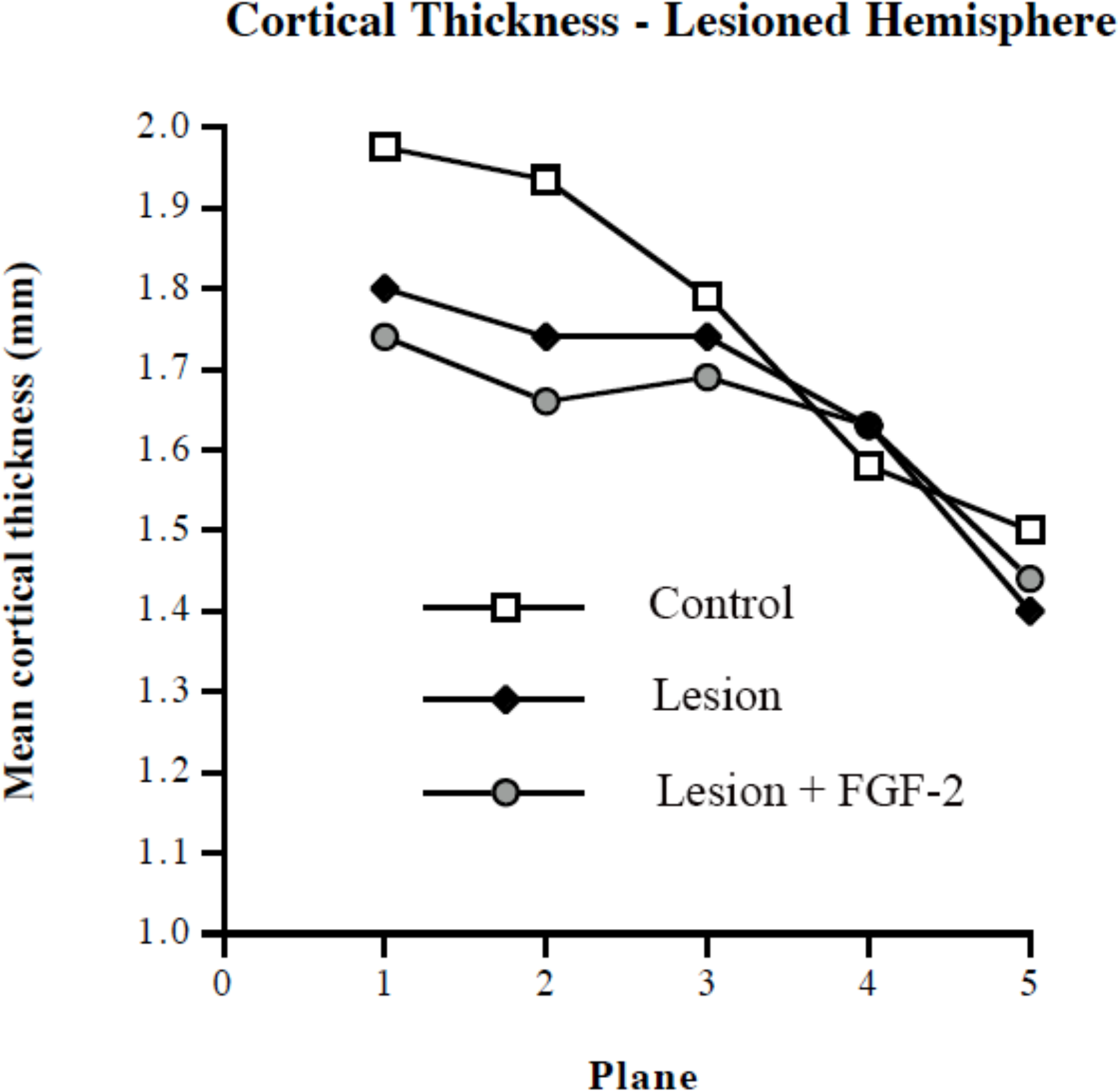
Mean cortical thickness (mm) in the lesion hemisphere in each of five planes. Because there was no treatment effect, data are collapsed across treatments.

One-way ANOVAs were done on each plane for each hemisphere. On the lesion side, both the lesion and lesion+ FGF-2 groups had significantly thinner cortices than controls in planes 1 and 2 (*p* < .05). Post hoc testing showed a significantly thinner cortex in the NT groups compared to the RT groups mainly due to the thinner cortex in the NT lesion+FGF-2 group in plane 3. On the intact side, the only significant difference was in the lesion+FGF-2 Complex housed group having a larger cortical thickness than controls, and the lesion RT group having a smaller cortical thickness than the corresponding CH and NT groups in plane 1. In summary, lesion and lesion+FGF-2 groups had overall thinner cortices in the lesion hemisphere, whereas in the intact hemisphere, there was a thicker cortex in the lesion+FGF-2 CH group.

### Dendritic analysis

The Golgi staining was good and comparable to that illustrated in many of our previous papers (e.g., Gonzalez & Kolb, 2003) Overall, complex housing produced a bilateral increase in dendritic length in both control and lesion animals whereas the lesion produced a bilateral decrease in dendritic length (Figure 4). Neither reach training nor FGF-2 affected dendritic length in the cells that were measured. One surprising result of the current study was that the lesion effect was essentially identical in the lesion and nonlesion hemispheres. The effects of complex housing and lesion were similar in the apical and basilar fields, so the ANOVA was done on the total length.

**Figure 4.**
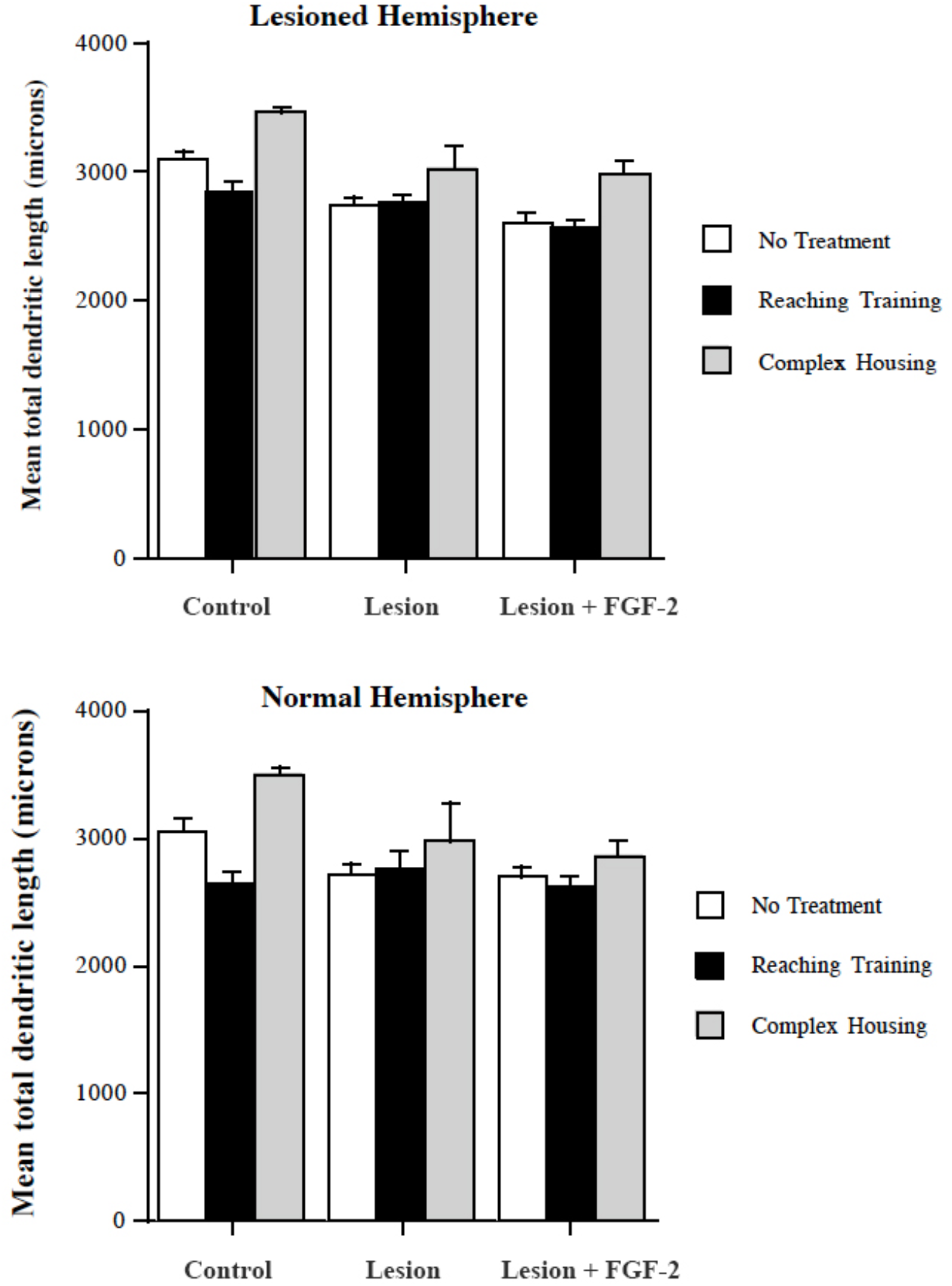
Summary of total dendritic length (apical plus basilar fields) measures in the lesion (top) and normal (bottom) hemispheres. Numbers represent means and SE.

ANOVA on the lesion hemisphere showed a main effect of lesion group, *F*(2, 95) = 19.4, *p* < .0001, and treatment, *F*(2, 95) = 21.8, *p* < .0001, but not the interaction, *F*(4, 95) = 0.84, *p* = .51. Post hoc tests (Fisher’s PLSD) on the lesion effect showed that the lesion and lesion+FGF-2 groups differed from control but not from one another. Similarly, on the treatment effect, the complex housed groups differed from the reach trained and no treatment groups that did not differ from one another. ANOVA on the normal hemisphere was essentially identical to the lesion hemisphere. There was a main effect of lesion group, *F*(2, 95) = 7.5, *p* = .0017, and treatment *F*(2, 95) = 9.3, *p* = .0005, but not the interaction, *F*(4, 95) = 0.84, *p* = .51. The post hoc tests showed the same differences as on the lesion side.

## Behavioral Results

### Skilled Reaching

All rats with motor cortex lesions were impaired at reaching for food. Animals in the reach training (RT) groups tended to stop reaching after approximately 30-45 min of daily training. The overall effect of treatment was that the complex housing (CH) improved skilled reaching as did the combination of training and FGF-2 (Figure 5). Neither training nor FGF-2 alone proved to be beneficial. There was little improvement after the first test day so the treatment effects appeared to be in the first three postoperative weeks. A repeated measures ANOVA with lesion group, training, and test days as factors revealed a significant main effect of group, *F*(2, 45) = 13.51, *p* < .0001, and training, *F*(2, 45) = 6.87, *p* = .0025, but not test day, *F*(4, 180) = .48, *p* = .76. The only significant interaction was time x treatment, *F*(8, 180) = 2.59, *p* = .01. Three distinct groupings appeared by test day five, performing from best to worst, respectively: (a) control CH and RT animals, (b) untreated controls, CH lesioned, CH lesion+FGF-2, and CH lesioned animals, and (c) untreated lesioned, untreated lesion+FGF-2, and RT lesioned animals. A two-way ANOVA on test day five with lesion group and training as factors showed a significant main effect of group, *F*(2, 45) = 9.94, *p =* .0003, and training, *F*(2, 45) = 5.85, *p* = .006, but no interaction, *F*(4, 45) = 1.15, *p* = .35. Follow-up post hoc tests revealed that the CH lesion, RT lesion+FGF-2, and CH lesion+FGF-2 groups performed as well as untrained controls.

**Figure 5.**
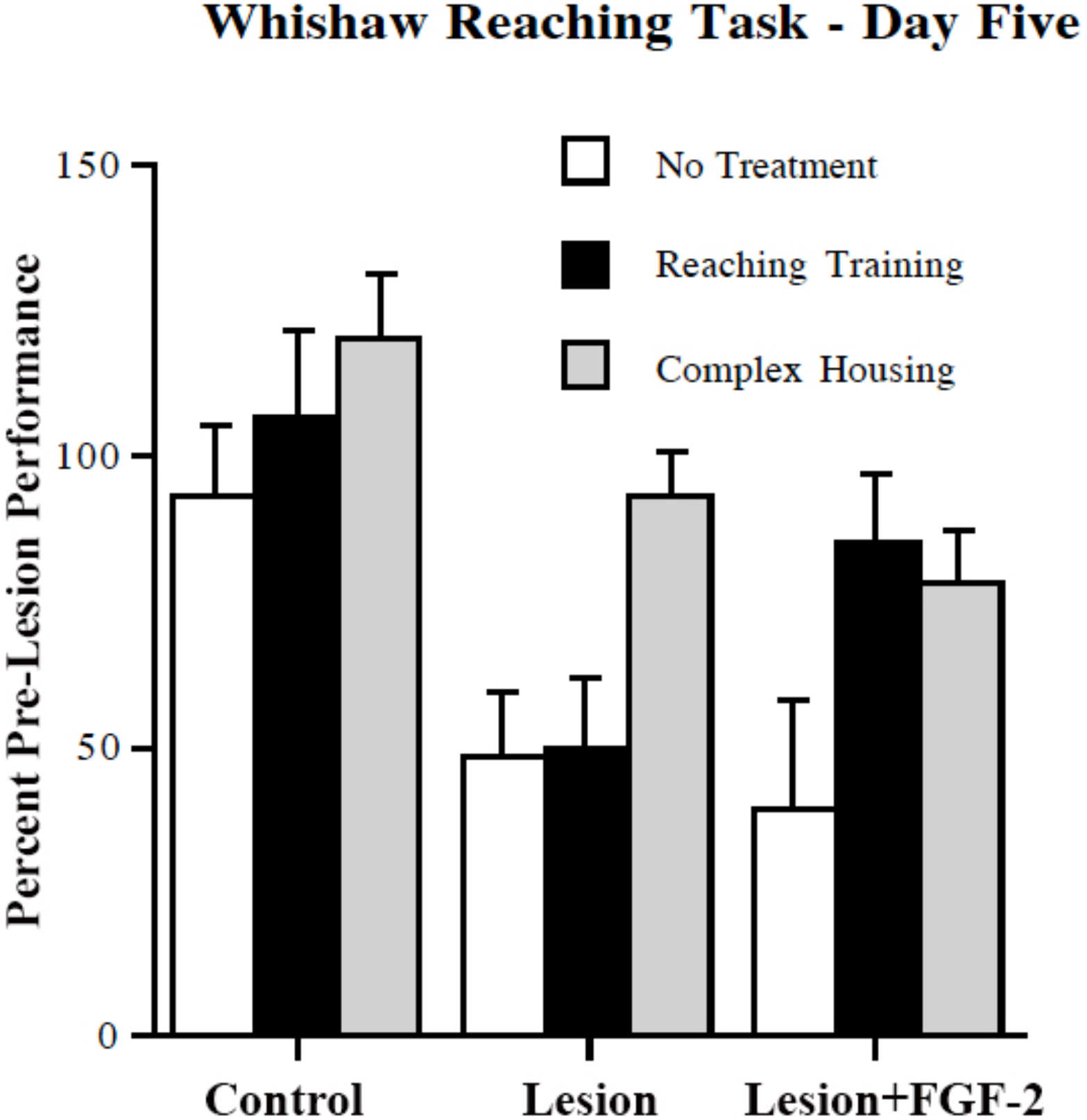
Percent accuracy on the Whishaw reaching task. A. Performance across the five test days, which began on poststroke day 21 and then were about 21 days apart. B. Summary of performance on test day five. Error bars represent SE.

### Spontaneous Vertical Exploration

Initially, animals actively explored the Plexiglas cylinder. As testing proceeded, animals tended to become habituated to the environment within the cylinder, and exploration decreased. They were more motivated to explore if they were food-deprived, and the scent of vanilla extract or chocolate chip cookie mash was near the top of the cylinder. Several of the CH animals even jumped to the rim of the cylinder. Over time, all trained groups decreased the use of their non-dominant paw relative to untrained groups, indicating an improvement in function of the affected paw (Figure 6). In other words, with training, animals were more likely to use their affected limb, or to increase the use of bilateral movements. A repeated measures ANOVA with lesion group, training, and test days as factors revealed a significant effect of group, *F*(2, 46) = 15.83, *p* < .0001, and training, *F*(2, 46) = 3.51, *p* = .038), but not test day, *F*(4, 184) = 1.63, *p* = .17). A two-way ANOVA on test day five with lesion group and training as factors showed an effect of group, *F*(2, 46) = 5.50, *p* = .007), and training, *F*(2, 46) = 3.29, *p* = .046), but no interaction, *F*(4, 46) = .97, *p* = .435) (Figure 6). The treatment effect was due to complex housing: Animals in the complex housing groups used their non-dominant paw significantly less often than did those in the treatment groups (Figure 6 bottom).

**Figure 6.**
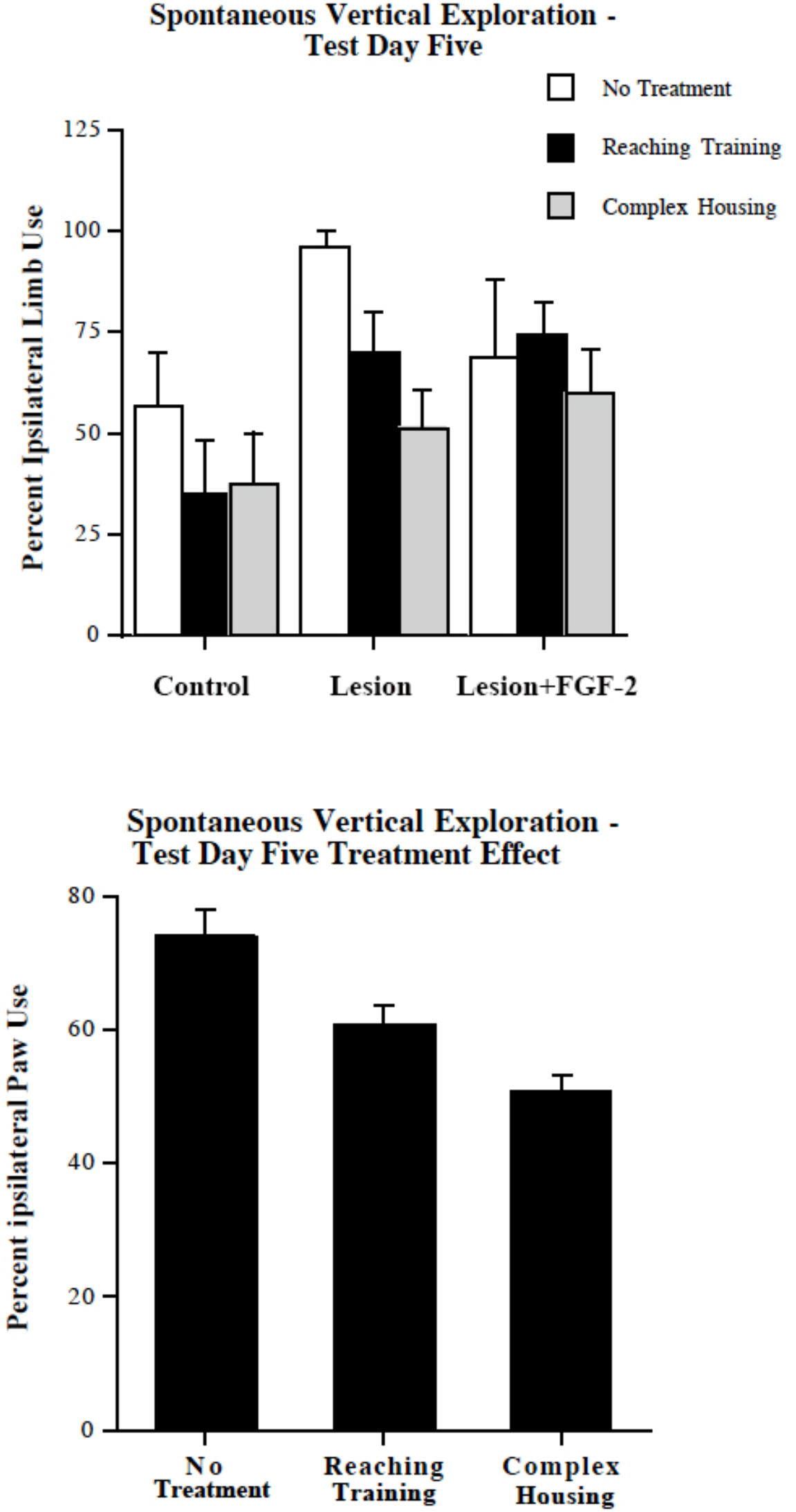
Top. Percent non-dominant wall leaning on the Schallert spontaneous vertical exploration task. A. Performance across the five test days. B. Summary of performance on test day five. Functional lesion and lesion+FGF-2 groups performed as well as the untrained controls. Bottom. Group effect without lesion effect. Error bars represent SE

### Forepaw Use During Swimming

After rats were trained to swim directly to the platform, control animals tended to hold both forelimbs motionless underneath the chin, whereas lesioned animals often used the impaired dominant limb to stroke (Figure 7). A repeated measures ANOVA with lesion group, training, and test days as factors revealed a significant effect of group, *F*(2, 46) = 13.28, *p* < .0001, no effect of training, *F*(2, 46) = .003, *p* =.997, but an effect of test day, *F*(4, 184) = 2.38, *p* = .054. The major effect of test day was mainly due to improvements in lesion+FGF-2 groups (Figure 7). A two-factor ANOVA on test day five with lesion group and training as factors revealed a group effect, *F*(2, 46) = 8.72, *p* = .0006, but no training effect, *F*(2, 46) =.16, *p* = .854) nor interaction, *F*(4, 46) = .54, *p* = .7101. Post hoc testing, however, indicated a significant effect comparing control and lesion, control and lesion+FGF-2, and lesion and lesion+FGF-2 groups. Thus, the lesion+FGF-2 groups displayed forepaw immobility in swimming more often than the other lesion groups (Figure 7 bottom).

**Figure 7.**
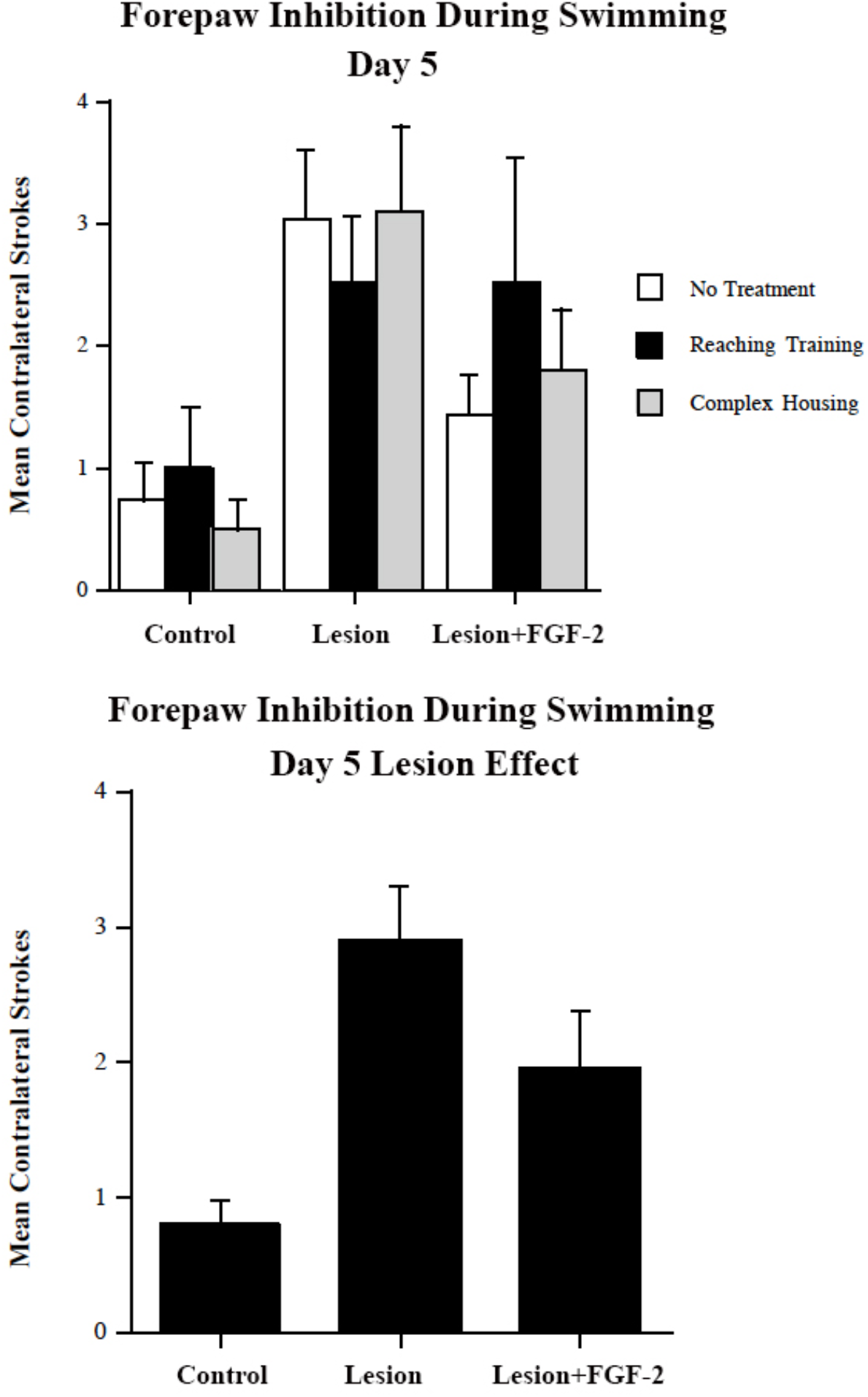
Top. Forepaw inhibition index on test day five. Note that FGF NT and CH groups performed best out of all lesion groups. Bottom. Group effect of forepaw inhibition during swimming on test day five. Error bars represent SE.

### Tongue Extension

Animals in all groups had tongue extension lengths of 9-13 mm, indicating the absence of damage in the lateral motor cortex, an area controlling tongue movements. ANOVA of mean tongue extension length collapsed over five test days indicated no effect of group, *F*(2, 45) = .024, *p* = .976, no effect of training, *F*(2, 45) = 1.82, *p* = .173, but an interaction, *F*(4, 45) = 5.42, *p* = .001. This interaction had to do with the complex housing lesion+FGF-2 group having a mean increase in tongue extension of about 1 mm.

### Single Pellet Reaching

Qualitative features of reaching were examined with this test to determine whether experience, lesion, and/or FGF-2 chronically altered the way the animals reached for food. Although animals were pre-trained to reach in the reaching cages, this specific task was new to them, and they thus needed training in this task for about 10 days before videotaping. Some lesion animals were unable to successfully use their dominant paw, and they were excluded from the analysis.

When rats reach for food, there is a stereotyped series of movements that can be analyzed into discrete components. Thus, a limb is lifted and positioned along the midline of the body by movements of the upper arm, with the paw located just below the mouth, so that it is aimed at the food. The head lifts to allow advance of the limb towards the food and as the limb travels the digits open. When the paw reaches the food, it is pronated so that the palm descends on the food and the digits grasp the food. During withdrawal, the paw is supinated at the wrist so that the food is presented to the mouth. As described by Whishaw et al. (1991), rats with motor cortex lesions have a severe deficit in pronating the paw over the food and in supinating the paw at the wrist to orient the food to the mouth. The latter deficit results in the rat placing the paw on the floor and then lowering the head to grasp the food with the teeth.

An analysis of the video films data was accomplished by examining each of 11 components separately. The principal finding was that FGF-2 treatment in combination with either tray reach training or complex housing reversed the supination deficit. These animals were now able to supinate the wrist and bring the food to the mouth. Curiously, the reach training produced a new deficit in the lesion alone animals as they were impaired at closing the digits to grasp the food. The food pellets in the single pellet task were about 1.5X larger than those in the tray reaching task so it is possible that the animals were unable to adjust to the larger pellet size. Thus, the specific training appeared to make the animals less flexible in learning a new motor task. Statistical analyses on each of the 11 components found that the FGF-2 plus either reach training or complex housing groups were significantly better than untreated lesion rats at *digits close, supination II*, and *release* (*p <* .05 or better).

### Claw Cutting

After animals were perfused, hindpaw nail length was measured to examine chewing and biting abilities required for the delicate movements of nail trimming (Whishaw et al., 1983). Mean nail length was calculated for each group with a mean length of about 4.5 mm for all groups except the FGF-2 +complex housing lesion group. A two-factor ANOVA revealed no effect of group, *F*(2, 43) = .68, *p* = .514, no effect of training, *F*(2, 43) = 1.65, *p* = .205, but an interaction, *F*(4, 43) = 5.88, *p* = .0007. The interaction was due to the beneficial effects of FGF-2 in the lesion and complex housing group.

## Discussion

This experiment compared the effects of a structured rehabilitation regime (skilled reaching) and more varied training (complex environment) with and without postlesion infusion of FGF-2 in rats having unilateral motor cortex lesions. Table 1 summarizes the results of testing in this experiment by listing the battery of tests administered and indicating whether there were significant functional improvements (+) across groups.

**Table 1.**
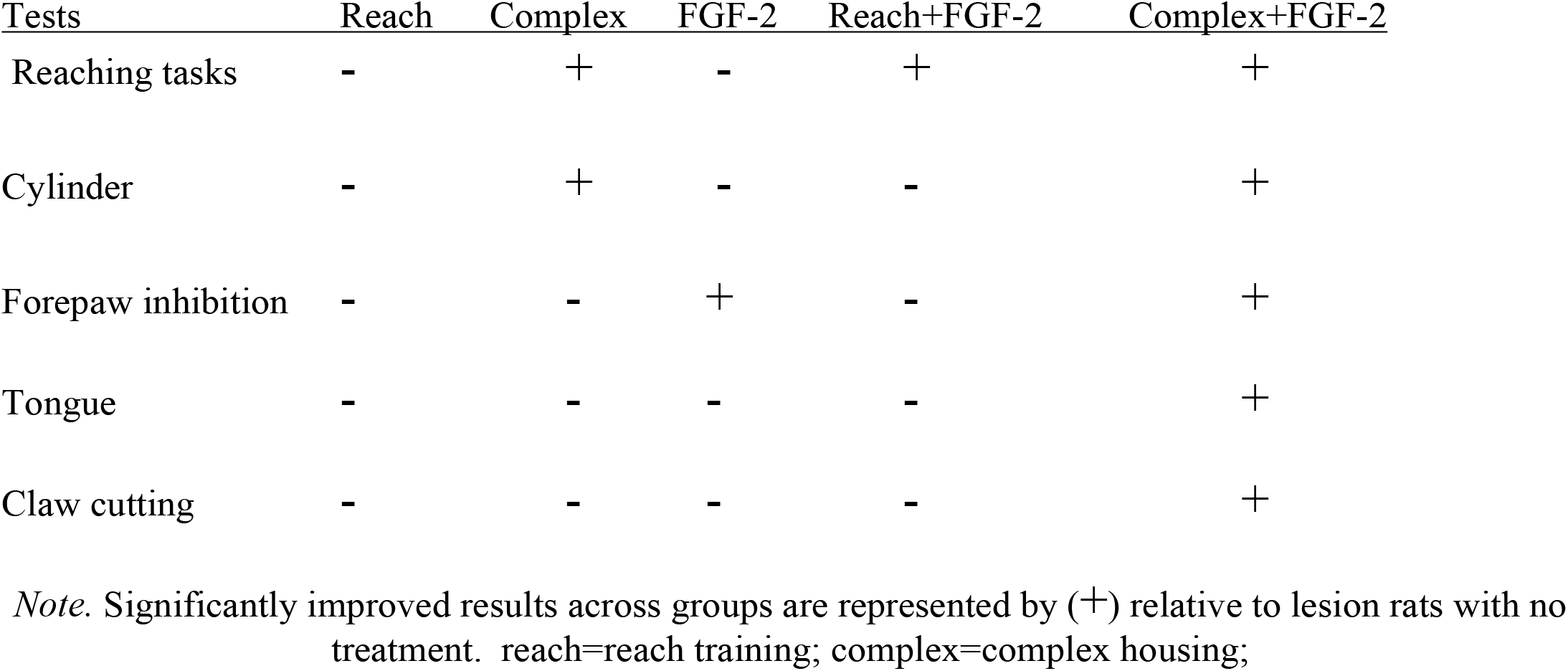
Comparison on a Battery of Tests Between Training/Treatment Groups.

| Tests | Reach | Complex | FGF-2 | Reach+FGF-2 | Complex+FGF-2 |
| --- | --- | --- | --- | --- | --- |
| Reaching tasks | - | + | - | + | + |
| Cylinder | - | + | - | - | + |
| Forepaw inhibition | - | - | + | - | + |
| Tongue | - | - | - | - | + |
| Claw cutting | - | - | - | - | + |

*Note.* Significantly improved results across groups are represented by (+) relative to lesion rats with no treatment. reach=reach training; complex=complex housing;

Overall, there were five principal findings from this study: (1) Most of the spontaneous recovery and the treatment effects occurred within the first three postoperative weeks; (2) There is a difference in rehabilitation-induced recovery of function between repetitive structured training and the behaviorally more diverse complex housing paradigms; (3) FGF-2 does enhance functional recovery, but only in combination with behavioral therapy; (4) There were bilateral dendritic changes related both to lesion and complex housing; and (5) Functional improvement in the FGF-2 plus behavior groups persisted for several months after the FGF-2 ended. Each finding will be discussed in turn.

### Behavioral Improved Occurred Over the First Three Weeks

In contrast to our previous studies (e.g., Gonzalez & Kolb, 2003; Whishaw et al., 2008) we did not begin our postoperative behavioral testing until three weeks after stroke. There were virtually no behavioral improvements after the first test whether or not animals had behavioral and/or FGF-2 treatments. Previous studies have found that improvements in skilled reaching after similar strokes are complete by 21 days poststroke and from the current study it appears that the same is true for the other measures used in the current study as well. Although it is difficult to extrapolate the comparable timeframe directly to humans, the results do suggest that poststroke treatments in humans are likely to be most effective in the immediate poststroke period, a conclusion also suggested by Biernaskie et al (2004).

### Reach-Training Versus Complex-Housing as Rehabilitation Regimes

Structured daily training alone over 4 months did not significantly improve rehabilitation-induced recovery of function on any subtest, unless FGF-2 was an additional treatment. Furthermore, the daily training on the same task interfered with performance of the lesion animals on the single pellet task. Complex housing dramatically improved performance on the tray reaching task but did not influence the qualitative performance on the single pellet task. There is considerable evidence that complex housing can produce enhanced functional outcome from a variety of forms of experimental brain damage including cortical ablation, cortical ischemia, and head trauma (e.g., Biernaskie & Corbett, 2001; Duran-Carabali et al., 2022; Johansson, 1996; Kolb & Elliott, 1987; Will & Kelche, 1992). Although the mechanism of the beneficial effects of complex housing is not known, it has been hypothesized that the treatment may increase the synthesis of neurotrophic factors, which in turn facilitate synaptic plasticity (e.g., Johansson, 2000; Cramer et al., 2011). Motor training has been shown to upregulate trophic factors such as BDNF and FGF-2 (see review by Kleim et al., 2003) so we had anticipated that the reach training would provide some benefit. One possibility is that although the training does enhance trophic factor production, the structured training produces synaptic changes that are too focal to provide much benefit after brain injury. It is known, for example, that although reach training does produce focal changes in synaptic motor cortex in otherwise normal rat brains, these changes are largely restricted to forelimb motor areas (e.g., Withers & Greenough, 1989). It seems likely that more widespread changes are necessary to facilitate functional recovery after motor cortical injury.

One unexpected finding was that reach training interfered with the performance of the single pellet task in the brain-injured group. One explanation was that the extensive training altered some aspect of behavior, such as posture, in such a way that the lesion animals were at a disadvantage in the new motor task. In fact, we have reported elsewhere that when animals are trained in the tray reaching task while on a low dose of nicotine they are impaired at learning the single pellet task later while off the drug and the reason appears to be that the animals have an abnormal posture that interferes with the learning of the new task (Gonzalez, Gharbawoe, Whishaw, & Kolb, 2005).

The failure to find beneficial effects of the repetitive training was surprising given that others have found some benefit (e.g., Dawson, Howarth, Tarnopolsky, Wong, & Gibala, 2003; Nudo, Wise, SiFuentes, & Milliken, 1996) and this type of training is often used by physiotherapists. It is possible that the absence of benefit in the current study is related to the task used. For example, Dawson et al. used a staircase task (Montoya et al., 1991) that is more difficult than the tray reaching used in the current study. Task difficulty thus may be critical for the benefits of repetitive structured training.

### FGF-2 Enhances Rehabilitation-Induced Recovery of Function

FGF-2 administration previously has been reported to enhance motor recovery after cerebral infarction (Kawamata et al., 1997) whereas in the current study there was no significant benefit. There are two important methodological differences, however. First, the infarcts were much more extensive in the Kawamato study, and they were produced by occlusion of the middle cerebral artery rather than by suction. The lesion etiology is known to influence the spontaneous synaptic reorganization following cortical injury and that may interact with neurotrophic factor treatment (Gonzalez & Kolb, 2004). Second, Kawamato et al. administered the FGF-2 intracisternally twice weekly for 4 weeks whereas we administered the FGF-2 intraventriculary with continuous infusion via minipump for 7 days. It is possible that the route and timing of administration is important, but this remains to be studied.

The combination of FGF-2 and either reach-training or complex housing proved to be functionally beneficial, although it was most helpful in the animals with complex housing. One reason for the beneficial effects of FGF-2 and behavioral therapy together is that the behavioral therapy was stimulating synaptic reorganization, which was in turn facilitated by the FGF-2. The added benefit of FGF-2 and complex housing may have resulted from the more extensive synaptic plasticity seen in animals in complex environments. Indeed, complex housing produces changes in most cortical regions, striatum, and hippocampus, all of which could potentially contribute to the observed functional enhancement. Curiously, the FGF-2 did not enhance the dendritic changes that we measured in parietal cortex, although there may have been changes in other forebrain areas that we did not measure. Indeed, the key finding here was that the combination of FGF-2 and complex housing enhanced function on every behavioral measure, suggesting that there was likely a widespread effect on the brain.

### Dendritic Changes Related to Injury and Complex Housing

There was a bilateral *decrease* in dendritic length related to the lesions and a bilateral *increase* in dendritic length related to complex housing. The complex housing essentially reversed the dendritic loss in the lesion animals. Curiously, there was no effect of FGF-2 on the cells measured, even though there was a beneficial effect of the combined FGF-2 and complex housing.

Complex housing is well known to increase dendritic arborization across the cortical mantle (see reviews by Kolb, 2020; and Kolb & Whishaw, 1998) and has previously been shown to be associated with dendritic growth and recovery in cortically-injured animals (Biernaskie & Corbett, 2001; Johansson, 2004). The surprising effect in the current study was the bilateral decrease in dendritic length in parietal cortex. We have shown previously that devascularizing lesions produce a bilateral increase in dendritic length in the motor and anterior cingulate cortex whereas aspiration lesions produce a bilateral decrease in cingulate cortex (Gonzalez & Kolb, 2003) and parietal cortex (Rowntree & Kolb, 1997). The decrease in parietal cortex with the aspiration lesions is thus consistent with our previous aspiration results. Similarly, several groups (e.g., Adkins, Voorhies, & Jones, 2004; Biernaskie & Corbett, 2001; Jones & Schallert, 1992, 1994) have shown increased dendritic length in undamaged motor cortex of animals with electrolytic, ischemic, or endothelin-1 lesions. The etiology of the injury thus seems to play a significant role in the endogenous changes in cortical organization as aspiration lesions appear to produce a different pattern of spontaneous dendritic changes than other forms of injury (see also Forgie, Gibb, & Kolb, 1996; Napieralski, Banks, & Chesselet, 1996, 1998). The fundamental difference between aspiration lesions and other forms of injury is that there is little dying tissue left in the brain. The differential effect of etiology on spontaneous changes in the brain must certainly interact with treatments and may account for why the FGF-2 and reach training were ineffective in the current study. The difference between the neural response to aspiration and other forms of cortical injury may also have implications for treatments after surgical excisions of cortical tissue such as in the treatment of epilepsy.

### Recovery of Function Over Time and Maintained Improvement

By testing animals repeatedly over the four-month recovery period we were able to demonstrate that there was little spontaneous recovery on the measures used. In addition, we showed that the benefits of complex housing or reach training and a week of FGF-2 treatment persisted for about 3.5 months after the FGF-2 treatment ended. Importantly, the benefits of either FGF-2 in combination with behavioral therapy or the complex housing alone was visible one month after the treatment began and was stable from then on.

The timing of onset of administration of the FGF-2 was immediate in the current study and behavioral training began either immediately (complex housing) or 3 days after injury (reach training). Parallel studies by others have shown that earlier initiation of rehabilitation is more beneficial than later administration (e.g., Barbay et al., 2001; Biernaskie et al., 2004) and we have shown elsewhere that housing animals in complex environments 3 months after perinatal injury provides little benefit (Comeau et al., 2005). It thus seems likely that the early administration of FGF-2 is critical to the functional benefits. It is equally important to note, however, that the early administration did not produce any obvious negative effects. It is known that forcing animals to use their affected limbs in the first days after injury can increase infarct volume (e.g., Schallert, Kozlowski, Humm, & Cooke, 1997) although less severe treatments can be beneficial (Bury & Jones, 2002; 2004). No differences in infarct volume or brain weight were present in the current study.

